# Epigenetic Dynamics in Meniscus Cell Migration and its Zonal Dependency in Response to Inflammatory Conditions: Implications for Regeneration Strategies

**DOI:** 10.1101/2024.07.22.604178

**Authors:** Yize Zhang, Yujia Zhang, Catherine Wang, Yuna Heo, Bat-Ider Tumenbayar, Se-Hwan Lee, Yongho Bae, Su Chin Heo

## Abstract

Meniscus injuries pose significant challenges in clinical settings, primarily due to the intrinsic heterogeneity of the tissue and the limited efficacy of current treatments. Endogenous cell migration is crucial for the healing process, yet the regulatory mechanisms of meniscus cell migration and its zonal dependency within the meniscus are not fully understood. Thus, this study investigates the role of epigenetic mechanisms in governing meniscus cell migration under inflammatory conditions, with a focus on their implications for injury healing and regeneration. Here, we discovered that a proinflammatory cytokine, TNF-α treatment significantly impedes the migration speed of inner meniscus cells, while outer meniscus cells are unaffected, underscoring a zonal-dependent response within the meniscus. Our analysis identified distinct histone modification patterns and chromatin dynamics between inner and outer meniscus cells during migration, highlighting the necessity to consider these zonal-dependent properties in devising repair strategies. Specifically, we found that TNF-α differentially influences histone modifications, particularly H3K27me3, between the two cell types. Transcriptome analysis further revealed that TNF-α treatment induces substantial gene expression changes, with inner meniscus cells exhibiting more pronounced alterations than outer cells. Gene cluster analysis pointed to distinct responses in chromatin remodeling, extracellular matrix assembly, and wound healing processes between the zonal cell populations. Moreover, we identified potential therapeutic targets by employing existing epigenetic drugs, GSKJ4 (a histone demethylase inhibitor) and C646 (a histone acetyltransferase inhibitor), to successfully restore the migration speed of inner meniscus cells under inflammatory conditions. This highlights their potential utility in treating meniscus tear injuries. Overall, our findings elucidate the intricate interplay between epigenetic mechanisms and meniscus cell migration, along with its meniscus zonal dependency. This study provides insights into potential targets for enhancing meniscus repair and regeneration, which may lead to improved clinical outcomes for patients with meniscus injuries and osteoarthritis.

## INTRODUCTION

The meniscus plays a crucial role in knee joint function, providing load-bearing support, shock absorption, and joint stability in the knee[1–3]. Prevalent among the middle-aged and elderly, meniscus injuries pose a considerable challenge, particularly within the avascular zone where intrinsic healing is limited[4–6]. Current treatment options often involve arthroscopic surgeries, including meniscectomy, which alleviate pain and prevent joint locking, but unfortunately, they many inadvertently hasten osteoarthritis progression[7–9]. Thus, elucidating the factors that govern meniscus repair and regeneration is crucial for devising effective therapies.

Pro-inflammatory cytokines, especially tumor necrosis factor alpha (TNF-α), are implicated in both meniscus injury[10] and osteoarthritis pathogenesis[11]. Studies have shown elevated TNF-α levels in synovial fluid post meniscus injury are associated with subsequent osteoarthritis[12–14]. Notably, TNF-α impedes the healing process of meniscus tissue post injury[11,15], whereas its inhibition via receptor antagonists fosters integrative meniscus repair[11]. Nonetheless, existing studies often overlook the zonal-dependent mechanical and biological properties of the meniscus in understanding the impact of injury on cellular behaviors in the meniscus.

The meniscus is composed of zones with distinct characteristics: the peripheral one-third (outer zone) is vascularized, whereas the inner one-third (inner zone) is avascular[16]. This zonal variation in vascularity likely influences cellular behavior and the response to injury. Additionally, the inner zone cells display a chondrogenic-like phenotype, in contrast to the fibroblastic nature of outer zone cells. Such phenotypic disparity leads to diverse extracellular matrix (ECM) compositions; the inner zone contains substantial amounts of both type-I and type-II collagen and higher aggrecan content, while the outer meniscus zone predominantly contains type-I collagen [16–19]. These zonal distinctions in ECM composition suggest that factors influencing meniscus repair and regeneration are zone dependent. Consequently, comprehending these zone-dependent differences is essential for developing effective repair and regeneration strategies.

Intrinsic healing of meniscus tears necessitates proper cell migration to the injury site, a process governed by intricate phenotypic and transcriptional changes, which are in turn tightly regulated by epigenetic modifications[20–22]. There is growing evidence that global histone reorganization, marked by alterations in chromatin condensation, is essential for efficient cell migration. These changes in chromatin structure facilitate the dynamic reconfiguration of the nucleus, thereby facilitating cellular movements during migration[22–24]. Advances in super-resolution microscopy such as Stochastic Optical Reconstruction Microscopy (STORM), have made it possible to visualize and quantify chromatin organization at the nanoscale[23,25,26]. Research employing STORM has demonstrated that a comprehensive reorganization of chromatin is necessary for meniscus cell migration within the dense fibrous networks of tissues[23]. However, the specific epigenetic mechanisms that drive the migration of meniscus cells, particularly the differential responses of inner and outer meniscus cells in response to injury and pro-inflammatory cytokines, remain to be elucidated. Tri-Methylation of Histone H3 lysine 27 (H3K27me3) is pivotal in orchestrating phenotypic and transcriptomic alterations, significantly impacting cell migration across diverse cell types[20–22]. The downregulation of H3K27me3 levels via siRNA-mediated knockdown of EZH2 (the enzyme responsible for this specific methylation as a subunit of the polycomb repressive complex 2) has been observed to modulate cell migration in fibroblasts, mesenchymal stem cells, epithelial cells, and cancer cells[27–30]. These observations highlight the regulatory importance of H3K27me3 in cell motility within various biological settings. However, the specific contributions of H3K27me3 to meniscus cell migration, particularly its role in cells from different zones of the meniscus before and after injury, remain to be fully elucidated.

Thus, the objective of this study is to investigate the impact and zonal dependency of epigenetic mechanisms on meniscus cell migration. We particularly aim to explore how inflammatory conditions modulate the migratory behaviors of inner and outer meniscus cells migration. Furthermore, we interrogate the epigenetic mechanisms by which inner and outer meniscus cells respond differently to proinflammatory cytokine TNF-α by focusing on discrepancies in histone modifications along with global chromatin reorganization. This research offers valuable insight into the intricate interplay between epigenetic factors, inflammation, and meniscus cell behavior. Notably, our findings reveal distinct histone modification patterns between inner and outer cells during migration, with significant variations in baseline H3K27me3 levels across meniscal zones. These differences underscore the regulatory role of H3K27me3 in meniscus cell migration. Additionally, our transcriptomic analysis and TNFR1 distribution suggest a heightened sensitivity of inner cells to TNF-α treatment. Furthermore, this study also highlights the therapeutic potential of epigenetic manipulation, as evidenced by the restoration of cell migration speed following the inhibition of specific histone modifiers.

Taken together, our study demonstrates that the migratory capacity of inner meniscus cells can be restored under TNF-α treatment through the small molecule modulation of histone methylation. This suggests a promising direction for epigenetic-targeted therapies aimed at intrinsic meniscus regeneration amidst inflammatory conditions. By advancing our comprehension of the epigenetic regulation of meniscus cell migration and its zonal dependency, this study makes a significant contribution to the field of meniscus repair and regeneration. Ultimately, these insights have the potential to enhance clinical outcomes for patients with meniscus injuries.

## RESULTS

### Differential Histone Modification Patterns and Chromatin Dynamics Drive Distinct Migration Behaviors in Inner and Outer Meniscus Cells

To assess the baseline migration behaviors of inner and outer meniscus cells, we conducted a wound closure “scratch” assay using meniscus cells harvested from juvenile bovine menisci and seeded onto 12 well tissue culture plates **(Figure 1a, b)**. Our analysis revealed that both inner and outer cells exhibited comparable migration speeds and wound closure rates at 6 and 12 hours post-scratch (**Figure 1c**).

**Figure 1:**
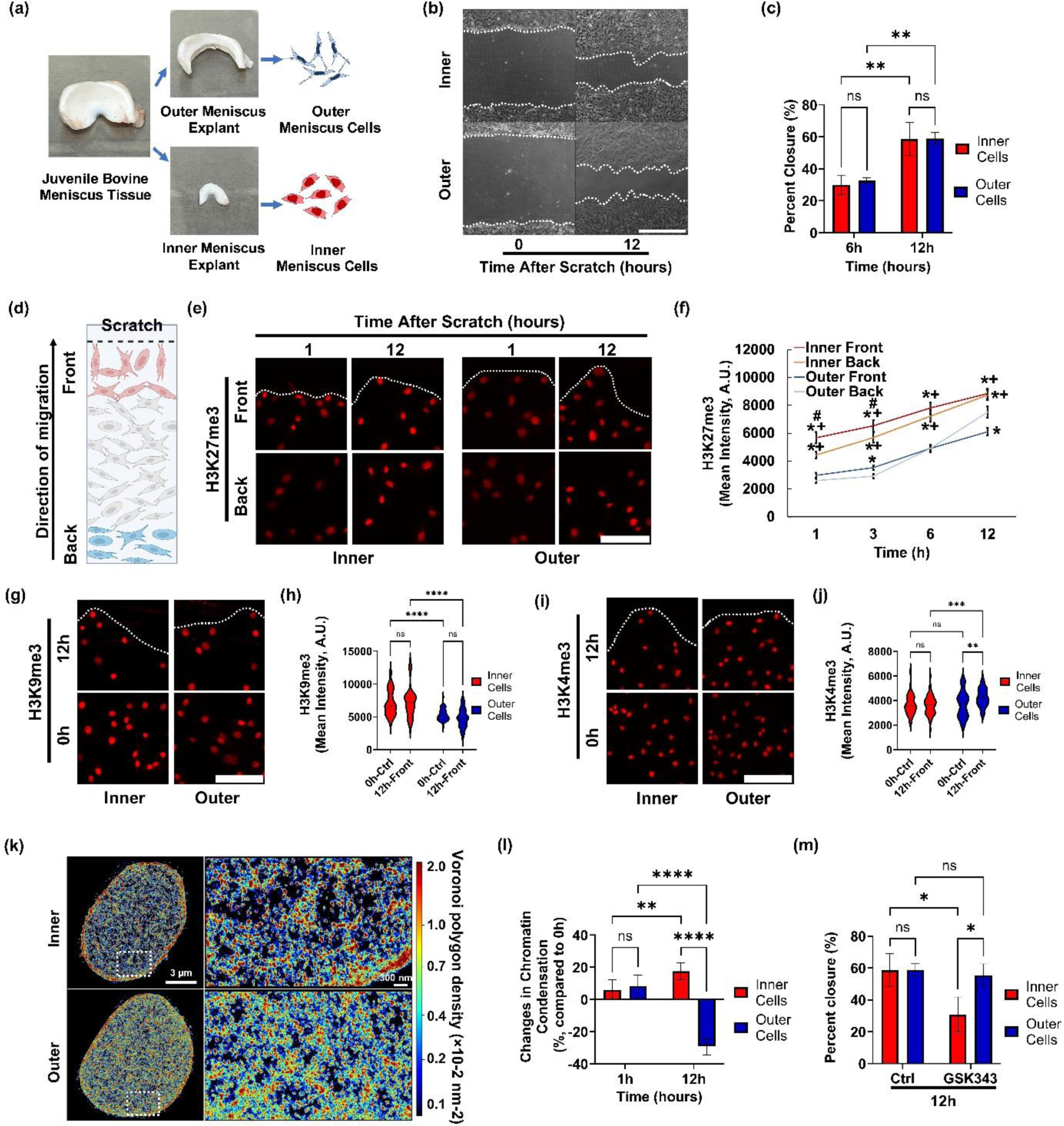
H3K27me3 requires for inner meniscus cell migration: (a) Schematic illustrating isolation of inner and outer zone meniscus cells. (b) Representative images of wound closure assay in inner or outer meniscus cells (scale bar: 500 µm) and (c) results quantified as percent wound closure over time (3 different donors in triplicates, **: p<0.01. Mean ± SD). (d) Schematic illustrating H3K27me3 immunofluorescence intensity measurement in inner or outer cells located at the migration front (Front) vs. 10 cell layers behind the migration front (Back). Representative immunofluorescence images and quantifications of (e, f) H3K27me3 (*: p<0.05 vs. Outer Back, +: p<0.05 vs. Outer Front, #: p<0.05 vs. Inner Back, Mean ± SD), (g, h) H3K9me3, or (i, j) H3K4me3 (scale bar: 50 µm) after scratch (n>100 cells/group, **: p<0.01, ***: p<0.001, ****: p<0.0001. Mean ± SD). (k) Representative H2B-STORM images (scale bar: 5 µm) and (l) percent change in chromatin condensation in nuclei at the migration “front” during migration (n=7/group, **: p<0.01, ***: p<0.001, ****: p<0.0001. Mean ± SD). (m) Comparison of inner vs. outer meniscus vell migration with/without treatment with an EZH2 Inhibitor (GSK343, 10 μM, n=6/group *: p<0.05, Mean ± SD).

Next, to explore the changes in histone modification landscapes during migration, we compared histone modification markers between cells at the leading edge (“Front”) and those at the rear back (“Back”) of the migration process (**Figure 1d**). This comparison aimed to discern histone modification differences between the cells actively migrating at the front and those lagging at the back. Notably, we observed that the intensity of H3K27me3 (Tri-methylation of lysine 27 on histone H3, a marker of repressed facultative heterochromatin) in inner cells at the migration "Front" was higher than that in inner cells at the migration "Back" one-hour post-scratch (**Figure 1d-f**). As migration progressed, the intensity of H3K27me3 gradually increased, with cells further from the migration front eventually matching the intensity levels observed at the migration leading edge over the course of 12 hours, suggesting dynamic shifts in histone modification patterns during migration (**Figure 1e, f**). More intriguingly, the baseline level of H3K27me3 in the inner cells was consistently higher than that in the outer cells throughout the migration (**Figure 1e, f**). Similarly, inner cells exhibited a higher average intensity of the heterochromatin marker H3K9me3 (Tri-methylation of lysine 9 on histone H3, a repressed constitutive heterochromatin marker), although no significant changes in H3K9me3 levels were detected in either cell type during migration (**Figure 1g, h).** There was no significant difference in the baseline intensity of the euchromatin marker H3K4me3 (tri-methylation of lysine 4 on histone H3, indicative of active euchromatin) between these cells (p>0.05), with an observed increase in intensity only in the outer cells during migration (**Figure 1i, j**). Similar changes were observed in H3K9ac (acetylation of lysine 9 on histone H3, an active euchromatin marker) throughout the migration process (**Supplementary Figure 1a**). These results point to distinct histone modification pattern alterations between inner and outer meniscus cell migration. Notably, H3K27me3 appears to play a more pronounced role in the migration of inner meniscus cells.

Upon noting the distinct roles of H3K27me3 in the migration of inner and outer meniscus cells, we further investigated differences in the global chromatin condensation level changes between these cell types during migration. Utilizing Super-resolution Stochastic Optical Reconstruction Microscopy (STORM), we examined the nanoscale organization of histone H2B (H2B) to assess the global chromatin condensation in the nuclei of both cell types. For this, the inner and outer meniscus cells were fixed at short-term (1 hour) and long-term (12 hours) intervals post-scratch to trigger migration. Initial observations at 1-hour post-scratch indicated increased chromatin condensation levels at the migration front in both inner and outer cells, suggesting that chromatin condensation is an early event in cell migration for both cell types (**Figure 1k, l**). However, interestingly, long-term observations at 12 hours revealed sustained chromatin condensation in inner cells, whereas outer cells exhibited a significant reduction in chromatin condensation compared to both control and inner cells (**Figure 1k, l**). This indicates a differential regulation of chromatin dynamics during extended migration periods in these cell populations.

Building on the observed changes in histone methylation during the cell migration, we further investigated the necessity of H3K27me3 (the histone modification with significant changes during migration and marked differences between inner and outer cells) for proper cell migration. For this, we utilized GSK343, a selective inhibitor of Enhancer of Zeste Homolog 2 (EZH2), which catalyzes the methylation of lysine 27 on histone H3. GSK343 was administered to both inner and outer cells approximately 12 hours prior to creating the scratch, and its presence was maintained during migration induction. The impact of EZH2 inhibition on cell migration was assessed using the wound closure assay (WCA). Intriguingly, while GSK343 treatment reduced the migration speed of inner meniscus cells, it did not affect the migration speed of outer meniscus cell (**Figure 1m**). These findings imply that H3K27me3 and its associated chromatin condensation play a more specific role in the regulation of inner meniscus cell migration.

### Zone-Dependent Effects of Inflammation on Meniscus Cell Migration via Nanoscale Histone Reorganization

Given the known inflammatory responses triggered by meniscal injury[12–14], which influence migratory behavior of endogenous meniscus cells [11,15], we further investigated the impact of TNF-α, a key pro-inflammatory cytokine, on the migration of meniscus cells. For this, juvenile bovine meniscus cells from the inner and outer zones were cultured in 12-well cell culture plates to 80% confluency before the pre-treatment with TNF-α overnight. Subsequently, a wound closure assay (WCA) was performed, and cell migration was visually assessed 12 hours post-induction (**Figure 2a**). Notably, TNF-α treatment markedly reduced the migration speed of exclusively the inner zone cells, whereas the outer zone cells’ migration remained unaffected when compared to the untreated control groups (Ctrl) (**Figure 2b, c**). These findings suggest zone-dependent responses of meniscus cells in response to inflammatory stimuli.

**Figure 2:**
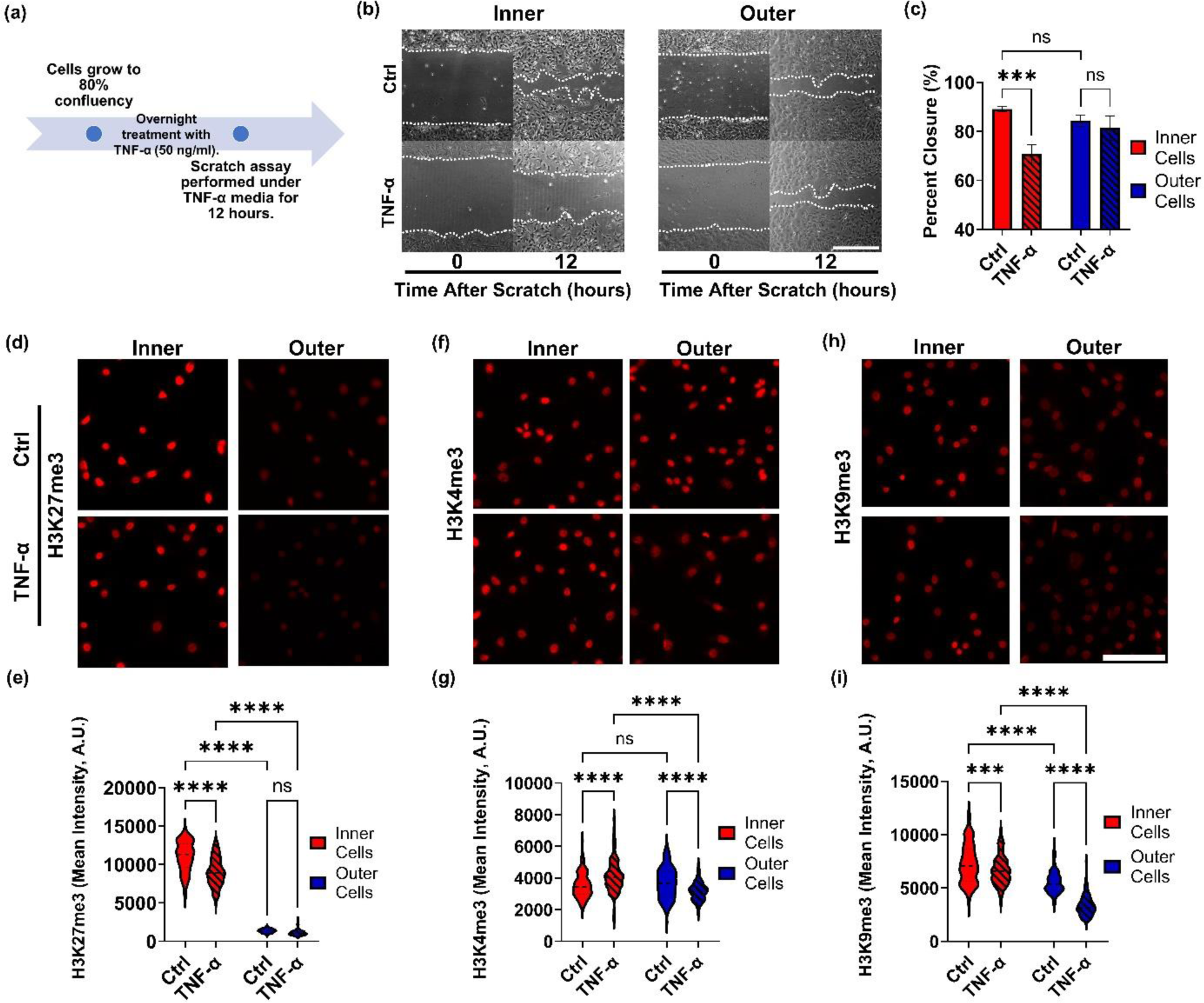
TNF-α decreases inner but not outer meniscus cell migration speed via modulation of H3K27me3 levels. (a) Schematic showing the design of the TNF-α treatment study (12-hour overnight treatment). (b) Representative images of wound closure assay (scale bar: 1 mm) and (c) results quantified as percent wound closure over time (performed in triplicate with cells from 3 different donors, ***: p<0.001, Mean ± SD). Representative immunofluorescence images and quantifications of (d, e) H3K27me3, (f, g) H3K9me3, or (h, i) H3K4me3 (scale bar = 100 µm) in inner or outer meniscus cells treated with/without TNF-α (***: p<0.001, ****: p<0.0001, n>100 cells/group).

Observing that histone modification levels fluctuate during meniscus cell migration (**Figure 1**), our subsequent analysis focused on the potential influence of TNF-α treatment on the migration velocity of meniscus cells via alterations in histone modification levels. To explore this, Inner and outer meniscus cells, cultured to 80% confluency, were subjected to fixation and staining for histone modification markers (H3K27me3, H3K9me3, and H3K4me3) following a 12-hour exposure to TNF-α. Consistent with prior observations in **Figure 1**, inner cells exhibited elevated baseline levels of the heterochromatin marker H3K27me3, which were notably diminished by TNF-α treatment—a response not mirrored in outer cells (**Figure 2d, e**). Conversely, the baseline levels of H3K9me3 were higher in inner cells, yet TNF-α treatment led to a reduction in both cell types, with outer cells experiencing a more pronounced decrease (**Figure 2h, i**). Interestingly, the euchromatin marker H3K4me3 displayed no initial disparity between cell types; however, TNF-α treatment resulted in an increase in inner cells and a decrease in outer cells (**Figure 2f, g**). These results suggest that TNF-α selectively modulates histone modification patterns in meniscus cells, particularly diminishing heterochromatin markers and augmenting euchromatin markers within inner cells. Further examination of global chromatin condensation levels during cell migration utilizing STORM-H2B imaging, revealed that inner cells at the migration forefront experienced a more substantial reduction in chromatin compaction following TNF-α treatment compared to outer cells (**Supplementary Figure 2**). These findings indicate an overall trend wherein inner cells respond more sensitively to TNF-α treatment regarding H3K27me3 levels and chromatin condensation status, thereby experiencing a more significant decrease in migration speed under inflammatory conditions. The variable effects of TNF-α on histone modifications underscore the zonal-dependent epigenetic mechanisms governing meniscus cell migration and their potential significance in the inflammatory sequelae of meniscal injuries.

### Transcriptomic Divergence in Inner and Outer Meniscus Cells in Response to TNF-α

Given the distinct impact of TNF-α on histone modifications and chromatin organization, influencing cell migration dynamics in inner and outer meniscus cells, we further hypothesized that TNF-α treatment would differentially alter the transcriptomic profiles in these cell populations. To explore this hypothesis, we performed a comprehensive whole-transcriptome analysis using mRNA samples from four experimental conditions: Outer control (OC), Outer TNF-α treatment (OT), Inner control (IC), and Inner TNF-α treatment (IT). By quantifying expression levels of 27,607 transcripts through next-generation sequencing, we aimed to uncover differences and commonalities across samples. Our unsupervised analysis, visualized via a Principal Component Analysis plot (PCA, **Supplementary Figure 3**) revealed that samples clustered primarily based on TNF-α treatment status along the PC1 axis, emphasizing the pivotal role of TNF-α in driving transcriptomic changes. Notably, within each treatment condition, the outer and inner meniscus samples exhibited distinct segregation along the PC2 axis, highlighting divergent transcriptomic responses to TNF-α treatment in these two cell subpopulations.

We further conducted a Likelihood Ratio Test (LRT) with a stringent threshold of p < 0.001 to identify 2,284 significantly altered transcripts across experimental conditions. To gain unbiased insight into the transcriptome alterations upon TNF-α treatment, we applied k-means clustering, an unsupervised machine learning method, on LRT-significant genes. K-means clustering divides data into k separate groups based on expression profile similarity, enabling us to identify different clusters with consistent gene expression patterns across samples without prior information. As shown in the heatmap (**Figure 3a**), the k-means clustering divided the 2,284 LRT-significant genes into six distinct clusters. Notably, clusters 1 and 5 exhibited divergent expression patterns in inner and outer meniscus cells following TNF-α treatment. To elucidate the functional relevance of these clusters, we performed Gene Ontology (GO) enrichment analysis. Cluster 1 was significantly enriched for biological processes related to chromatin remodeling, cell cycle regulation, and protein localization to chromosomes (**Figure 3b**). This suggests distinct responses of inner and outer meniscus cells in these cellular processes upon TNF-α treatment. Furthermore, cluster 5 was enriched for processes associated with extracellular matrix assembly, wound healing, and cell migration (**Figure 3c**). Focusing specifically on cell migration-related genes, TNF-α treatment elicited distinct responses in inner versus outer meniscus cells. Inner meniscus cells exhibited more pronounced changes in migration-related gene expression levels compared to outer cells, indicating heightened sensitivity to TNF-α treatment (**Figure 3d**).

**Figure 3.**
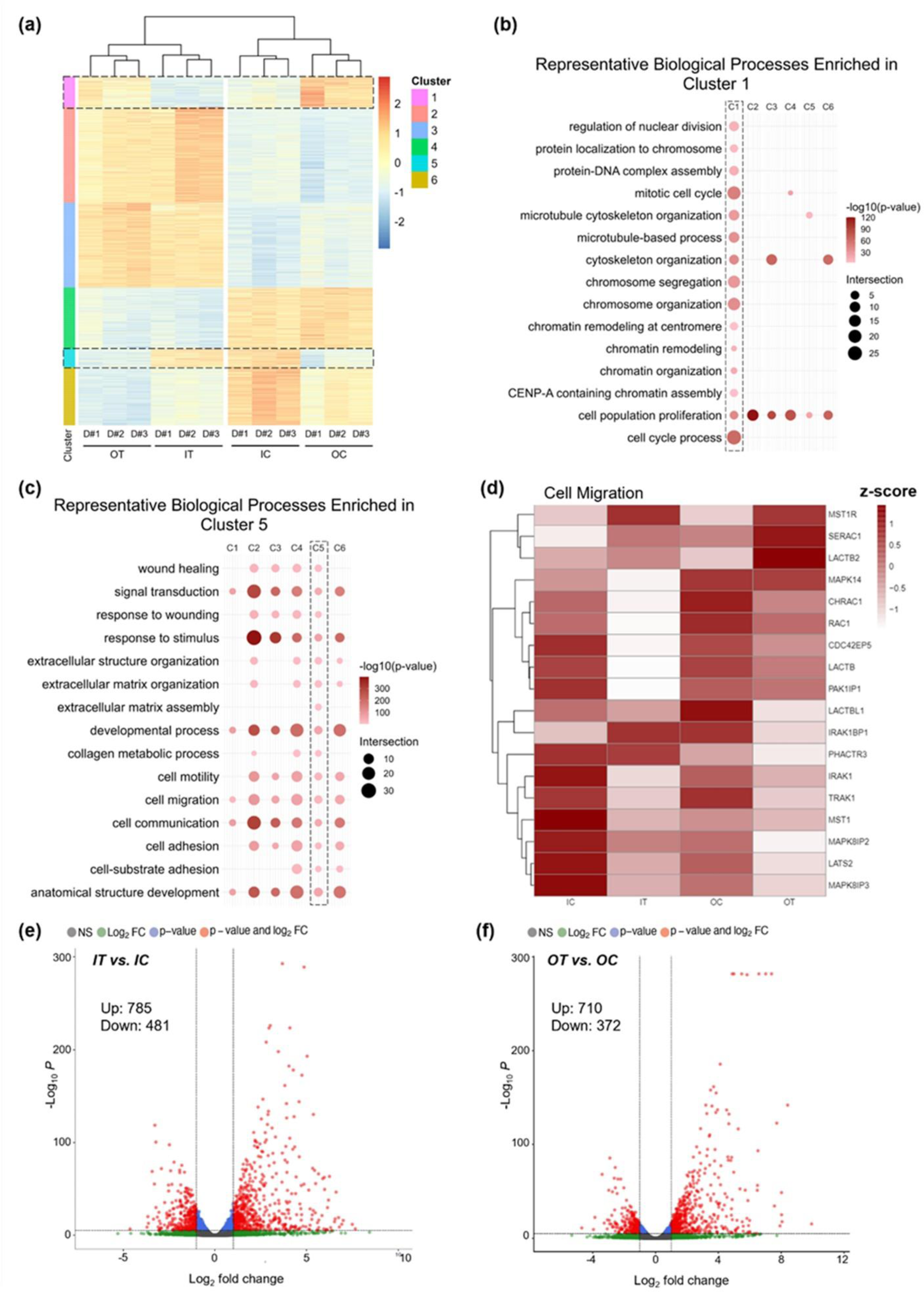
Transcriptomic landscape alterations in inner and outer meniscus cells upon TNF-α treatment. (a) Heatmap displaying expression patterns and K-means clustering (6 clusters) of 2,284 likelihood ratio test (LRT)-significant genes (p < 0.001) across four conditions: Outer Control (OC), Outer TNF-α (OT), Inner Control (IC), and Inner TNF-α (IT). Bubble plots showing representative biological processes enriched in (**b**) cluster 1, related to chromatin remodeling, cell cycle, and protein localization to chromosomes, and (**c**) cluster 5, associated with extracellular matrix assembly, wound healing, and cell migration. Bubble size represents the number of genes associated with each process and color intensity shows significance level-log_10_(p-value). (d) Heatmap displaying expression pattern of cell migration-related genes across IC, IT, OC, and OT measured in Z-score. Volcano plots show the distributions of differentially expressed genes (DEGs) in (**e**) IT vs. IC or (**f**) OT vs. OC comparisons.

To further investigate the differential transcriptomic responses elicited by TNF-α treatment in inner and outer meniscus cells, we performed differential expression analysis and identified differentially expressed genes (DEGs). This analysis revealed a total of 1,266 DEGs when comparing Inner control (IC) versus Inner TNF-α treatment (IT), with 785 genes upregulated and 481 genes downregulated (**Figure 3e**). Similarly, the comparison between Outer control (OC) and Outer TNF-α treatment (OT) identified 1,082 DEGs, consisting of 710 upregulated and 372 downregulated genes (**Figure 3f**). Notably, an examination of the top 30 upregulated and downregulated genes indicated that over 50% of these genes exhibited divergent expression between the inner and outer meniscus cells (**Supplementary Tables 1 and 2**). These data indicate the unique transcriptomic changes induced by TNF-α treatment in each cell population, reflecting their distinct responses to inflammatory conditions.

### Differential Expression and Distribution of TNF-α Receptors in Inner and Outer Meniscus Cells

The distinct migration behaviors of inner and outer meniscus cells under TNF-α treatments led us to hypothesize that these differences might be attributed from variations in the presence of TNF-α receptors in these cells. Thus, we focused on two major TNF-α receptors, TNF-α Receptor 1 (TNFR1) and TNF-α Receptor 2 (TNFR2), both integral to cellular processes in mammalian cells. Employing macroscopic immunofluorescence and super-resolution STORM imaging, we compared the baseline levels of TNFR1 and TNFR2 in both cell types. Immunofluorescence analysis indicated a significantly higher intensity of TNFR1 in inner meniscus cells compared to outer cells (**Figure 4a, b**). Additionally, at the nanoscale level, we employed STORM imaging for cells fixed and stained with TNFR1 antibodies and then calculated the density of TNFR1 present on the cell membrane. Consistent with the immunofluorescence data, STORM images of TNFR1 revealed a significantly higher density in inner cells compared to outer cells (**Figure 4c-e**). We also observed that TNFR2 expression was higher in inner compared to outer meniscus cells, though the differences are not as significant as those for TNFR1 (**Supplementary Figure 4**). These findings suggest that the differential distribution and density of TNF surface receptors between the inner and outer meniscus cells contribute to the divergent responsiveness to TNF-α. Specifically, the higher levels and density of TNFR1 in inner meniscus cells may lead to more significant changes in histone modification and chromatin condensation levels, thereby influencing cell migration under inflammatory conditions.

**Figure 4:**
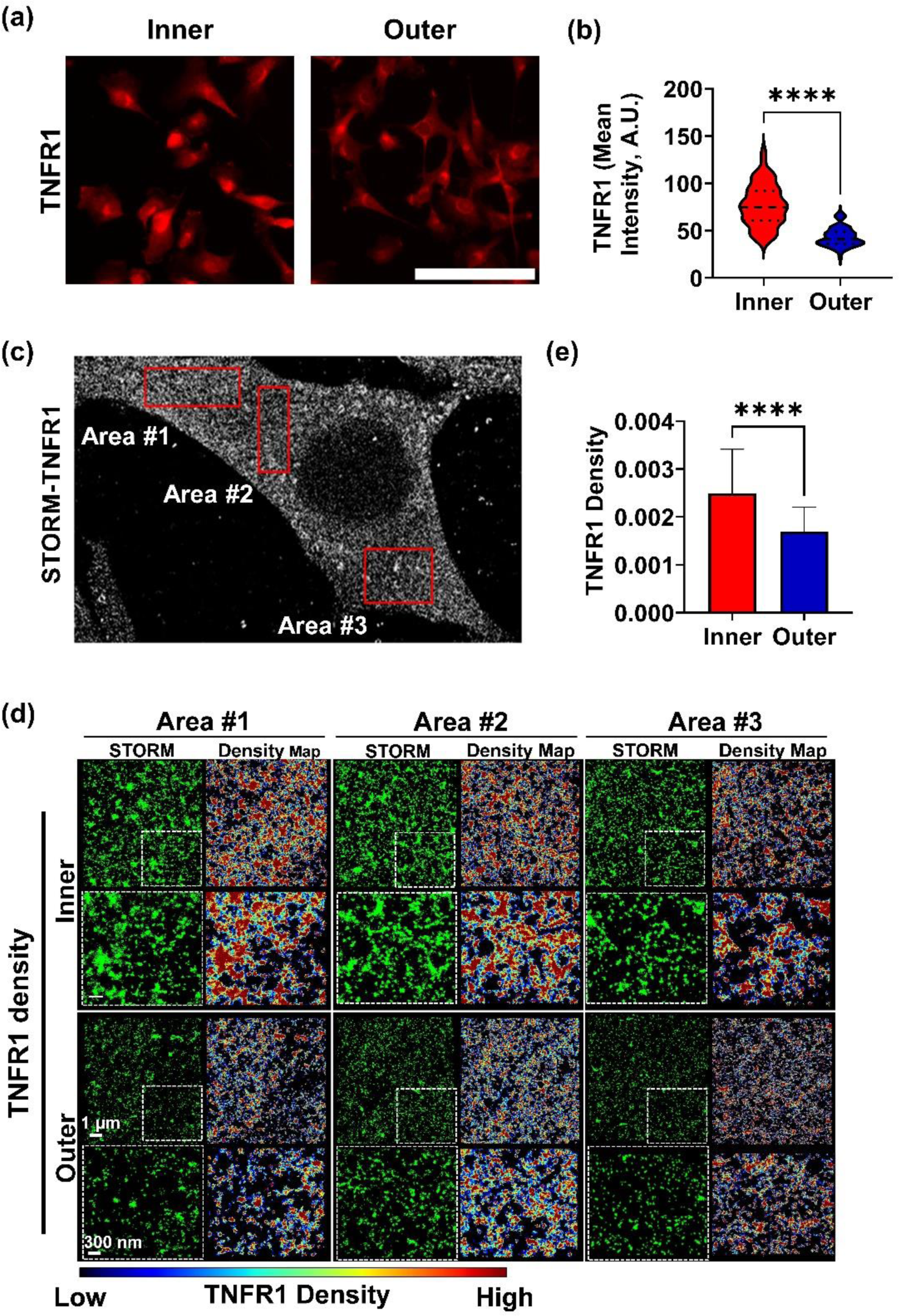
Baseline TNFR1 expression is higher in inner cells compared to outer cells. (a) Representative immunofluorescence images of TNF-α Receptor 1 (TNFR1) in inner or outer cells (scale bar: 100 μm) and (b) quantification of TNFR1 intensity (n>100 cells/group, ****: p<0.0001). (c) Nanoscale TNFR1 distribution measured in three random areas per cell, and representative (d) TNFR1 STORM images and density map, and (e) quantifications of TNFR1 density (number of TNFR1 localizations per total area, demonstrating significantly higher density in inner cells, n>100 measurements/group, Mean ± SD).

### Selective Inhibition of Histone Enzyme Activation Restores TNF-α-Mediated Migration in Inner Meniscus Cells

Recognizing the pivotal role of H3K27me3 in TNF-α-mediated reduction of inner meniscus cell migration speed, we next hypothesized that altering the histone methylation levels affected by TNF-α could potentially restore the impaired migration speed. To test this hypothesis in this study, we leveraged established epigenetic drugs targeting key histone modification processes. Specifically, we employed GSKJ4, an inhibitor of the H3K27me3 demethylase JMJD3, known to elevate the overall level of H3K27me3 in cells[31–34], and C646, an inhibitor of the histone acetyltransferase p300, which reduces the overall level of histone acetylation[35–37].

To evaluate the potential therapeutic strategies for meniscus healing, we evaluated the effects of GSKJ4 and C646 on the migration speed of inner meniscus cells under TNF-α-induced inflammatory conditions. Our wound closure assay results were promising, showing that a 12-hour treatment with GSKJ4 at both low (2 μM, GSKL) and high doses (5 μM, GSKH) partially restored the migration capacity of TNF-α-treated inner meniscus cells (Figure 5a-c). Similarly, pretreatment with C646 at low (10 μM, C646L) and high doses (30 μM, C646H) also led to partial recovery in migration capacity under TNF-α treatment (**Figure 5a-c**). These findings suggest that the epigenetic modulation by GSKJ4 and C646 can effectively counteract the histone methylation alterations induced by TNF-α, potentially fostering a transcriptional environment conducive to cell migration. Considering the chronic inflammation and impaired healing associated with meniscal injuries, repurposing GSKJ4 and C646 emerge as a novel therapeutic approach to enhance recovery. Future studies should focus on the long-term effects of these treatments on meniscus cell behavior and their potential to enhance functional recovery *in vitro* and *in vivo*. Further exploration of the gene expression profiles and signaling pathways affected by these epigenetic drugs will provide insights into their mechanisms of action for developing targeted therapies for musculoskeletal injuries. Taken together, this study reveals that the modulation of histone methylation and acetylation can significantly influence the migratory behavior of meniscus cells, potentially offering new therapeutic intervention for enhancing tissue repair.

**Figure 5:**
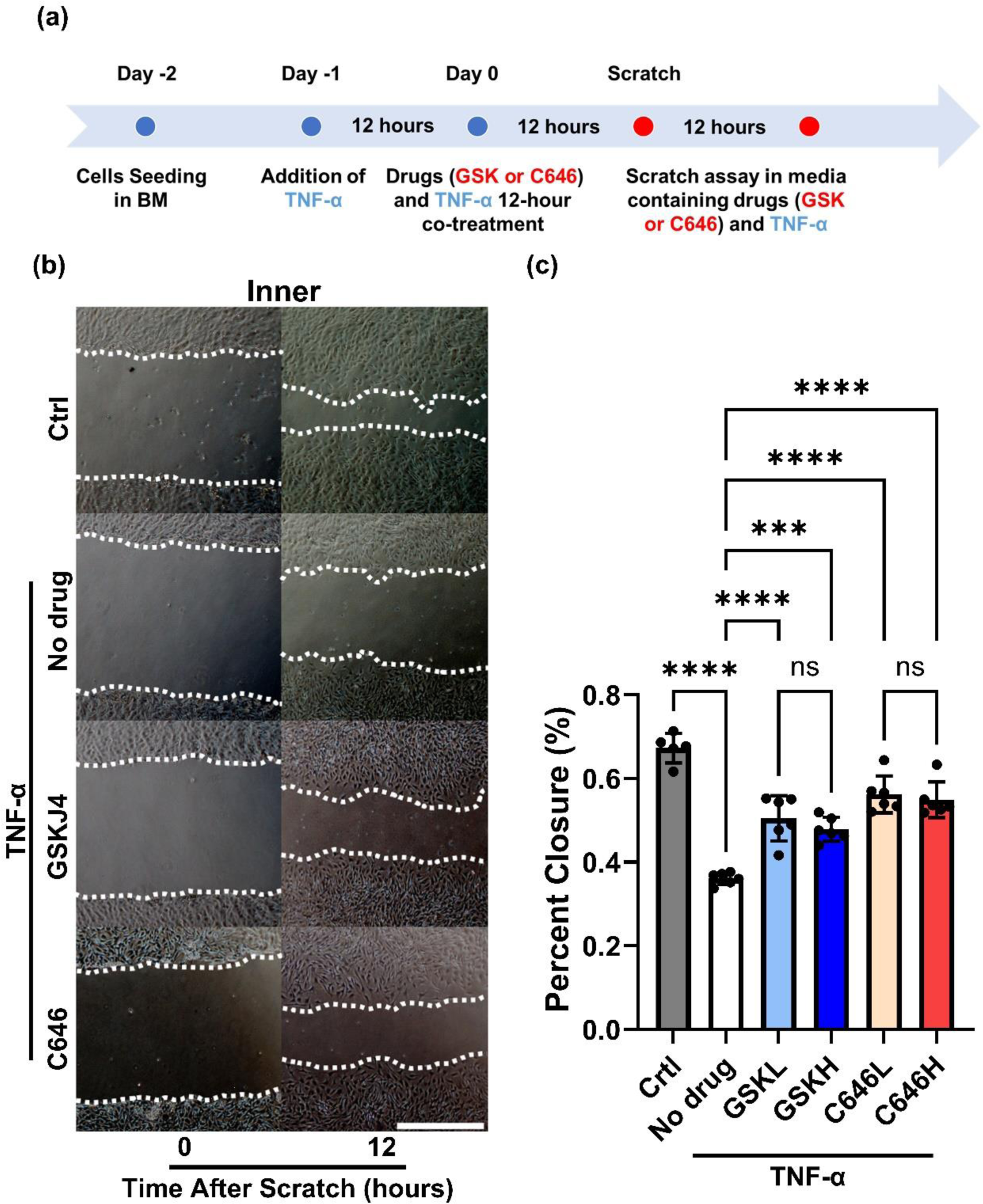
GSKJ4 and C646 Restore Inner Meniscus Migration Speed Under TNF-α Treatment. (a) Schematic illustrating the design of the epigenetic drug treatment study. Inner meniscus cells were subjected to TNF-α overnight treatment, followed by 12 hours of co-treatment with TNF-α and epigenetic drugs before performing the 12-hour scratch assay. (b) Wound closure assay images demonstrating the effects of drug treatments (scale bar = 1 mm). (c) Quantification of wound closure over time, expressed as percent wound closure (n = 5-6 per condition from 3 different donors; ***: p<0.001, ****: p<0.0001. Mean± SD, L: low dose, H: high dose).

## DISCUSSION

Meniscus injuries are common and challenging to treat, particularly in the avascular inner zone where current interventions like suturing or meniscectomy fall short. Thus, there is a pressing need for the development of new treatment methods. To address the unmet clinical needs, in this study, we aimed to investigate the impact of epigenetic mechanisms on meniscus cell migration under inflammatory conditions, and understand the differential histone modification patterns and chromatin dynamics driving distinct cell migration behaviors with the ultimate goal of pioneering epigenetic-based therapies for meniscus repair. Here, we established that the histone modification marker H3K27me3 plays a critical role in facilitating the proper migration of inner meniscus cells under inflammatory conditions. Furthermore, we elucidated the alterations in chromatin condensation and transcriptomic profiles that contribute to the observed migration pattern of inner meniscus cells.

Our objectives were to elucidate the epigenetic mechanisms influencing meniscus cell migration under inflammatory conditions and to explore therapeutic avenues for enhancing meniscus repair. To this end, we exposed inner and outer meniscus cells to TNF-α known to impede repair processes[38–40]. Notably, our experiments confirmed that TNF-α significantly curtails the migration speed of inner meniscus cells without affecting outer cells. This led us to probe deeper into the associated histone modification and transcriptomic changes. Consequently, we administered existing epigenetic drugs targeting histone acetyltransferase and histone demethylase to restore the migratory capability of inner meniscus cells under inflammatory conditions.

We discovered pronounced differences in histone modification patterns and chromatin organization between the two cell types during migration. Notably, inner meniscus cells exhibited higher levels of the heterochromatin marker H3K27me3 compared to outer meniscus cells during migration. The selective inhibition of EZH2, which aimed to reduce H3K27me3 levels, resulted in a decreased migration speed exclusively in inner cells, aligning with findings in other cell types[27–30]. Conversely, outer meniscus cells remained unresponsive to EZH2 inhibition, underscoring a specific regulatory role for H3K27me3 in inner cell migration. Furthermore, TNF-α treatment was found to selectively reduce H3K27me3 levels in inner cells. While the exact mechanism behind TNF-α’s induction of zonal-dependent epigenetic changes is not fully understood, the differential response hints at distinct sensitivities to pro-inflammatory signals between inner and outer meniscus cells. This could be due to the higher density of TNF-α receptors on inner meniscus cells. These insights accentuate the importance of considering the zonal properties of the meniscus when addressing its inflammatory response. The zonal-dependent impact of TNF-α on histone modifications could be pivotal in developing targeted therapies for meniscus injuries.

Our study elucidated differential changes in global chromatin condensation between inner and outer meniscus cells during extended migration. We observed that chromatin condensation in inner cells not only persisted but increased over time, whereas it decreased in outer cells. This observation supports the "pack and go" hypothesis, which posits that chromatin condensation is essential for initiating cell migration[22–24]. The sustained increase in chromatin condensation in inner cells may reflect enduring transcriptional changes linked to their migratory phenotype. Conversely, the reduction in chromatin condensation in outer cells could suggest a distinct transcriptional program that facilitates their migration. To understand the disparate epigenetic responses of inner and outer meniscus cells to TNF-α treatment, we analyzed the TNF surface receptors. Our findings indicate a higher abundance and density of TNF-α Receptor 1 (TNFR1) in inner cells, correlating with the reduced migration speed under inflammatory conditions. The pronounced presence of TNFR1 implies a heightened sensitivity of inner cells to TNF-α, leading to downstream epigenetic alterations, particularly affecting H3K27me3 levels, which are crucial for the migration of inner meniscus cells.

To explore the cellular mechanisms driving changes in cell migration induced by inflammation, we employed RNA sequencing (RNA-seq) to profile transcriptomic alterations in response to TNF-α. Principal component analysis (PCA) identified TNF-α as a significant influencer of transcriptomic shifts, demonstrating the divergent transcriptomic responses between inner and outer meniscus cell populations. Employing a stringent Likelihood Ratio Test (LRT) and k-means clustering, we identified 2,284 significantly altered transcripts and six distinct clusters with unique expression profiles. Notably, clusters pertinent to chromatin remodeling and cell migration displayed contrasting expression patterns between the two cell types. Interestingly, key cell migration-related genes such as MAPK14 and MAPK8IP3 were found to be downregulated in inner meniscus cells post-treatment, which are recognized for their role in promoting cell migration[41]. Additionally, genes associated with the Rho GTPase family, including Rac1 and CDC42EP5, exhibited reduced expression in inner cells, providing a potential mechanism underlying their diminished migration speed under inflammation conditions[42]. Additionally, the differential expression analysis revealed the distinct cellular mechanisms activated in each population. These findings suggest the importance of considering zonal-dependent properties in meniscus repair and regeneration. The variable regulation of chromatin dynamics emphasizes the meniscus’s inherent heterogeneity, calling for tailored approaches to cater to the specific cellular responses during the healing process. Through the elucidation of these molecular pathways, our goal is to forge more effective methods for facilitating meniscus repair and regeneration, ultimately enhancing clinical outcomes for patients with meniscus injuries.

A significant accomplishment of this study is the novel application of existing epigenetic drugs for treating meniscus tear injuries. Our selected drug candidates, GSKJ4 (a histone demethylase inhibitor)[31–34] and C646 (a histone acetyltransferase inhibitor)[35–37] effectively restored the migration speed of the inner meniscus cells under TNF-α treatment. This aligns with prior research suggesting that inflammatory cytokines activate JMJD3 via the NF-kB pathway, diminishing H3K27me3 levels[43]. Both GSKJ4 and C646 have demonstrated anti-inflammatory effects in various models, linking their roles in limiting inflammation with promoting cell migration in meniscus cells[44,45]. Similarly, C646 has shown promise in alleviating inflammatory responses in conditions such as inflammatory lung disease [46] and anti-bacterial macrophages[47].

Our study also marks the first to demonstrate that only inner meniscus cells modify key heterochromatin markers, particularly H3K27me3, in response to TNF-α, affecting their migration speed. While the exact mechanisms by which H3K27me3 alterations regulate gene expression related to cell migration are yet to be fully understood, future research employing epigenetic sequencing techniques such as ChIP-seq and ATAC-seq will be instrumental in uncovering the epigenetic landscape governing cell migration. Moreover, considering the significant differences in cell migration patterns between 3D and 2D environments, the use of advanced 3D migration assays will be crucial in further validating the efficacy of anti-TNF-α therapy in meniscus tissue engineering.

In conclusion, this study enhances our understanding of the interplay between epigenetic mechanisms, zonal-dependent properties, and meniscus cell migration. Our findings also highlight the importance of considering meniscus heterogeneity and provide insights into potential targets for improving meniscus repair and regeneration. These results suggest promising avenues for developing targeted interventions to enhance clinical outcomes for individuals with meniscus injuries and osteoarthritis. Further research is needed to elucidate the specific molecular mechanisms underlying the observed epigenetic changes and their functional consequences in meniscus repair and regeneration. The approach employed in this study also shows promise for treating injuries in other tissues, including tendon, muscle, cardiovascular, skin, and neural tissues, warranting further exploration.

## MATERIAL & METHODS

### Meniscus cell isolation and culture

Juvenile bovine menisci (< 3 months, Research 87, Inc., Boylston, MA) were harvested and divided into the inner and outer meniscus areas (**Figure 1a**). The respective meniscus tissues were then cut into 5 mm cubes. To isolate meniscus cells[23,48], the inner and outer meniscus tissue cubes were cultured separately in basal growth media consisted of Dulbecco’s Modified Eagle’s medium supplemented with 10% fetal bovine serum and 1% penicillin, streptomycin, and fungizone (PSF) in an incubator maintained at 37°C and 5% CO_2_ for approximately 14 days, after which inner and outer meniscus cells were collected. To ensure consistent cell conditions, only passage 1 (P1) cells were used for all subsequent experiments.

### Wound closure assay

To assess 2D migration of inner and outer meniscus cells, a wound closure “scratch” assay (WCA) was conducted. Cells (at passage, P1) were seeded onto the twelve-well tissue culture dish (2×10^^5^ cells per well). The cells were cultured additionally for 1-2 days until they reached 90% confluency. Subsequently, the WCA was initiated by creating a scratch on the surface of the plate using a 20 μl standard pipette, forming one horizontal and one vertical scratch intersecting at the center of the plate. Images were captured using an Inverted Microscope (Nikon ECLIPSE TS100) at 0, 6, or 12 hours post-scratch. The percent closure of the wound was determined by analyzing the images with ImageJ.

For the drug or pro-inflammatory cytokine treatments during the WCA, P1 cells were initially seeded in the twelve-well tissue culture dishes at a density of 2×10^^5^ cells per well and cultured until reaching 80% confluency (typically taking 1-2 days). To investigate the effect of the EZH2 inhibitor on cell migration speed, cells were then treated with GSK343 (SML0766-Sigma-Aldrich) at concentrations of 10 μM in BM overnight before conducting the “scratch” assays. To examine how the inflammatory cytokine TNF-α influences meniscus cell migration speed, cells were treated with TNF-α (T7539-Sigma-Aldrich) at concentrations of 50 ng/ml prior to the “scratch” assays (**Figure 3a**).

To explore the potential of using existing small molecule histone modification drugs to restore inner meniscus cell migration speed under inflammatory conditions, cells were pretreated with TNF-α (T7539-Sigma-Aldrich) overnight before co-treatment with C646 (a histone acetyltransferase inhibitor, SML0002-Sigma Aldrich) at a concentration of 10 µM (Low dose, L) or 30 µM (High dose, H) or GSKJ4 (a histone demethyltransferase inhibitor, ab144395-Abcam) at concentration of 2 µM (L) or 5 µM (H) for 12 hours prior to the scratch assay (**Figure 6a**).

### RNA-Sequencing

To investigate the transcriptomic level changes leading to different responses to TNF-α between inner and outer meniscus cells, RNA sequencing was performed on both cell types with and without TNF-α treatment. Passage 1 (P1) cells were seeded in six-well tissue culture dishes at a density of 1×10^6^ cells per well. The cells were cultured for an additional 1 to 2 days until they reached 90% confluency. For the control group, total RNA from inner and outer meniscus cells was collected after 2 days of culture using TRIzol Reagent (Invitrogen). For the TNF-α treatment group, cells were treated with TNF-α (T7539-Sigma-Aldrich) at a concentration of 50 ng/ml in 3 ml of BM overnight (12 hours) before RNA collection using the same method. The collected RNA was purified using Direct-zol RNA Microprep Kits (Zymo Research). RNA sample quality was assessed with a NanoDrop Spectrophotometer (Thermo Fisher Scientific), a Qubit Fluorometer (Thermo Fisher Scientific), and an Agilent 2100 Bioanalyzer (Agilent Technologies). RNA-Seq library preparation (rRNA depletion) and sequencing (150 bp paired-end) were performed at Genewiz (South Plainfield, New Jersey). Sequence reads were trimmed to remove possible adapter sequences and low-quality nucleotides using Trimmomatic v.0.36. The trimmed reads were then mapped to the Bos taurus reference genome (available on Ensembl) using the STAR aligner v.2.5.2b. Unique gene hit counts were calculated using featureCounts from the Subread package v.1.5.2.

Differential expression analysis between the experimental conditions Outer Control (OC), Outer TNF-α (OT), Inner Control (IC), and Inner TNF-α (IT) was performed using the DESeq2 package of R programming language [49]. The EnhancedVolcano R package was used to visualize the global transcriptional change across the groups compared[42]. The DESeq2 Likelihood Ratio Test (LRT) was applied to test for differential expression between contrasts of interest. A stringent significance threshold of p < 0.001 was used for this analysis to ensure a robust level of confidence in the rejection of the null hypothesis. Additionally, the DESeq2 package was used to identify differentially expressed genes (DEGs). DEGs were selected based on the following thresholds: Benjamini-Hochberg adjusted p-value of < 0.05, absolute log2(fold-change) of > 1.0. R pheatmap package[50] was used to illustrate expression patterns, clusters, and distributions of LRT-significant genes.

To gain a deeper understanding of the functional profiling associated with LRT-significant genes, a gene enrichment analysis was performed to examine highly enriched biological processes. The analysis was carried out using the g:GOSt function on the gProfiler web server (https://biit.cs.ut.ee/gprofiler/gost)[51]. The significance threshold was set to the Benjamini-Hochberg FDR (false discovery rate) value, and significant Gene Ontology (GO) terms were defined by an adjusted p value of ≤0.05. For visualization purposes, bubble plots for the representative GO terms were generated using the ggplot2 package of R[52].

### Immunofluorescence

To examine the dynamics of various histone modification markers during cell migration and drug treatments, cells were fixed with 4% paraformaldehyde for 20 minutes at 37°C at different time points, specifically 1, 6, and 12 hours after creating the scratch. Cells were permeabilized for 5 minutes in a solution of 0.05% Triton X-100 in PBS supplemented with 320 mM sucrose and 6 mM magnesium chloride.

For fluorescence labeling of histone modification markers, cells were incubated at 4°C overnight with primary antibodies targeting H3K27me3 (Rabbit, 9733 CST, 1:300, Thermo), H3K9me3 (Mouse, 6F12 CST, 1:300, Thermo), and H3K4me3 (Rabbit, 9751 CST, 1:300, Thermo), H3K9ac (Rabbit, MA5-11195, 1:200, Thermo) in PBS. Cells were then incubated for 60 minutes at room temperature with Alexa-Fluor 546 goat anti-mouse or anti-rabbit secondary antibody (1:200; Thermo Fisher). Images were acquired using a widefield microscope (Leica DM6000) at 20× magnification and analyzed for fluorescence mean intensity at the migration front and migration back (**Figure 2a**) using ImageJ.

To assess the effects of TNF-α treatment on histone modification marker expressions in inner and outer meniscus cells, inner and outer cells were seeded on 24 well-treated plates for 1-2 days were then treated with 50 ng/mL TNF-α (T7539-Sigma-Aldrich) overnight (12 hours). Fixed cells were then stained and imaged for H3K27me3, H3K9me3, H3K4me3, or H3K9ac. Additionally, to investigate the distribution of the TNF-α surface receptors-1 or 2 (TNFR1 or TNFR2) on inner and outer meniscus cells, cells were fixed and stained with primary antibodies against TNFR1 (Rabbit, 1:200, ADI-CSA-815; ENZO) and TNFR2 (Rabbit, 1:200, ab15563; Abcam). Images were acquired using a widefield microscope (Leica DM6000) at 20× magnification and then analyzed for fluorescence mean intensity per cell using ImageJ.

### STORM imaging

To investigate the effect of TNF-α treatment on chromatin condensation levels at the nanoscale in the cells, inner and outer meniscus cells were seeded on 8-well cover glass chambers (Nunc™ Lab-Tek™ II Chambered Coverglass) for 2 days in BM before conducting the WCA. TNF-α (50 ng/ml) was added to the chamber, and cells were treated overnight with TNF-α, followed by fixation with methanol-ethanol (1:1) at -20 °C for 7 minutes. To optimize antibody staining and minimize nonspecific bindings, a blocking buffer (BlockAid, ThermoFisher) was added to each well and incubated for 1 hour. Diluted anti-Histone H2B (Rabbit, 1:50, 15857-1-AP; Proteintech) was then added to the wells and incubated overnight at 4°C to specifically label the H2B histone protein.

Following sample preparation, samples were washed and subsequently incubated for 40 minutes with secondary antibodies custom-labeled with activator-reporter dye pairs (Alexa Fluor 405–Alexa Fluor 647, Invitrogen) for STORM imaging[23,25,26,53]. All imaging experiments were conducted using a commercial STORM microscope system (ONI Nanoimager). During imaging, the 647-nm laser was utilized to excite the reporter dye (Alexa Fluor 647, Invitrogen) and switch it to the dark state. Subsequently, a 405-nm laser was employed to reactivate the Alexa Fluor 647 in an activator dye (Alexa Fluor 405)–facilitated manner. An imaging cycle was implemented in which one frame belonging to the activating light pulse (405 nm) was alternated with three frames belonging to the imaging light pulse (647 nm). To ensure proper photo-switching of the Alexa Fluor 647, an imaging buffer, prepared according to a previously established protocol [23,25,26,53], was utilized. The imaging buffer comprised 10 mM cysteamine MEA (Sigma Aldrich, 30070-50G), 0.5 mg/ml glucose oxidase, 40 mg/ml catalase (Sigma), and 10% glucose in PBS [23,25,26,53]. Cellular localization at the migration front was achieved using a 640 nm laser to excite the Alexa Fluor 647 dye, acquiring a total of 20,000 frames of images per nucleus. STORM image localizations were obtained using Nanoimager software (ONI) and rendered with custom-written software Insight3 (a gift from B. Huang, UCSF, USA). For quantitative analysis, custom-written MATLAB codes were employed [23,25,26,53]. TNFR1 nanoscale distribution was quantified via random sampling of three different locations on the cell surface (three locations covering approximately 50% of the cell area/cell), and then the ratio of localization/area (the number of localizations with detected localization/the total pixels in the sampled area) was analyzed (**Figure 4c**).

### Statistical Analysis

Statistical analysis was conducted using GraphPad Prism 9 (La Jolla, CA). For comparisons between two groups, a Student’s t-test was utilized, and two-way analysis of variance (ANOVA) with Tukey’s honestly significantly different (HSD) post hoc test was used for multiple group comparisons. Transcriptomic data analysis was performed using the DESeq2 Likelihood Ratio Test (LRT) to assess differential expression between contrasts of interest, applying a significance threshold of p < 0.001, Benjamini-Hochberg adjusted p-value of < 0.05, and absolute log2(fold-change) of > 1.0. The R pheatmap package was employed to visualize expression patterns, clusters, and distributions of LRT-significant genes, as well as the associated Gene Ontology terms. Results are presented as means ± SD, and differences were considered statistically significant at p < 0.05. Sample and replicate numbers are provided in the figure legends.

## Acknowledgments

This research was supported by grants from the National Institutes of Health (K01 AR07787, R01 AR079224, R01 HL163168) and the NSF Science and Technology Center for Engineering Mechanobiology (CMMI-1578571). We extend our gratitude to Dr. Robert L Mauck, Dr. Melike Lakadamyali, and Dr. Joel D. Boerckel for their invaluable scientific insights and contributions to this study.

## Author Contributions

The conception and design of the studies were collaboratively undertaken by all authors. The execution of biological experiments was carried out by Yize Zhang, Yujia Zhang, C. Wang, Y. Heo, and S. Lee. Yujia Zhang and B. Tumenbayar were instrumental in the analysis of RNA-seq data. All authors played a pivotal role in the analysis and interpretation of the data, as well as in the drafting and critical revision of the manuscript, culminating in the final submission.

## Supplementary Information

**Supplementary Figure 1.**
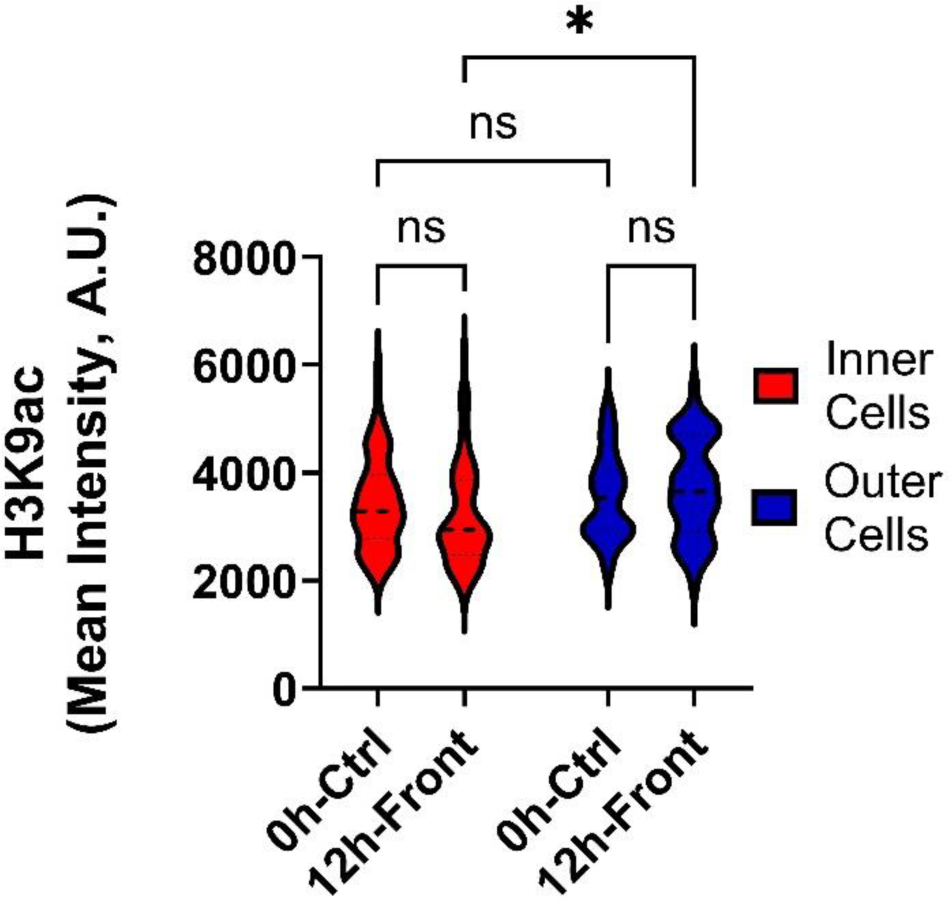
Changes in H3K9ac levels during inner and outer meniscus cell migration. Quantification of H3K9ac Immunofluorescence results for Inner and Outer meniscus cells, measured before and 12 hours after scratch (n > 50 cells/group). *: p<0.05, Mean± SD).

**Supplementary Figure 2.**
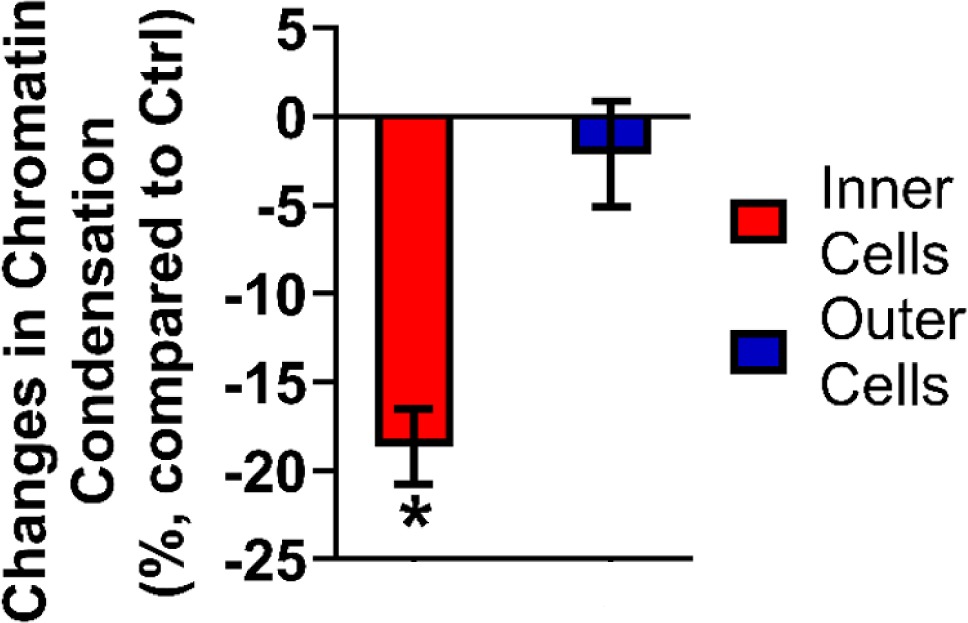
Global chromatin condensation changes for inner and outer meniscus cell migration under TNF-α treatment. Percent change in chromatin condensation in nuclei at the migration "front" for inner and outer meniscus cells after 12-hour TNF-α (50 ug/ml, n = 7/group, *: p<0.0001. Mean± SD).

**Supplementary Figure 3.**
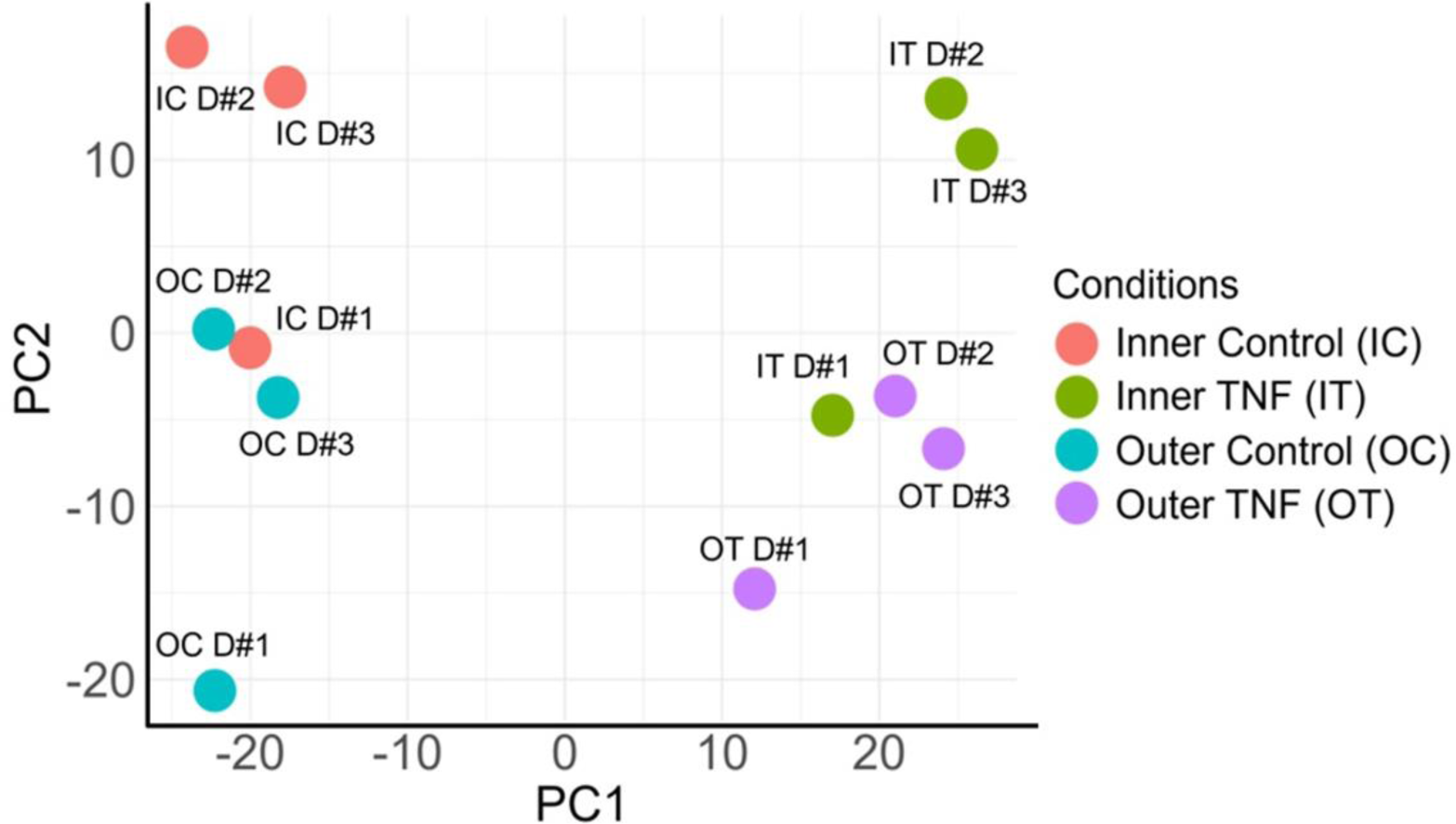
Global transcriptomic profiles of inner and outer meniscus cells. Principal Component Analysis (PCA) plot illustrating the distinctions and similarities between the four experimental conditions based on their global gene expression profiles. Each point represents a sample, highlighting the clustering patterns that differentiate inner and outer meniscus cells under varying treatments.

**Supplementary Figure 4.**
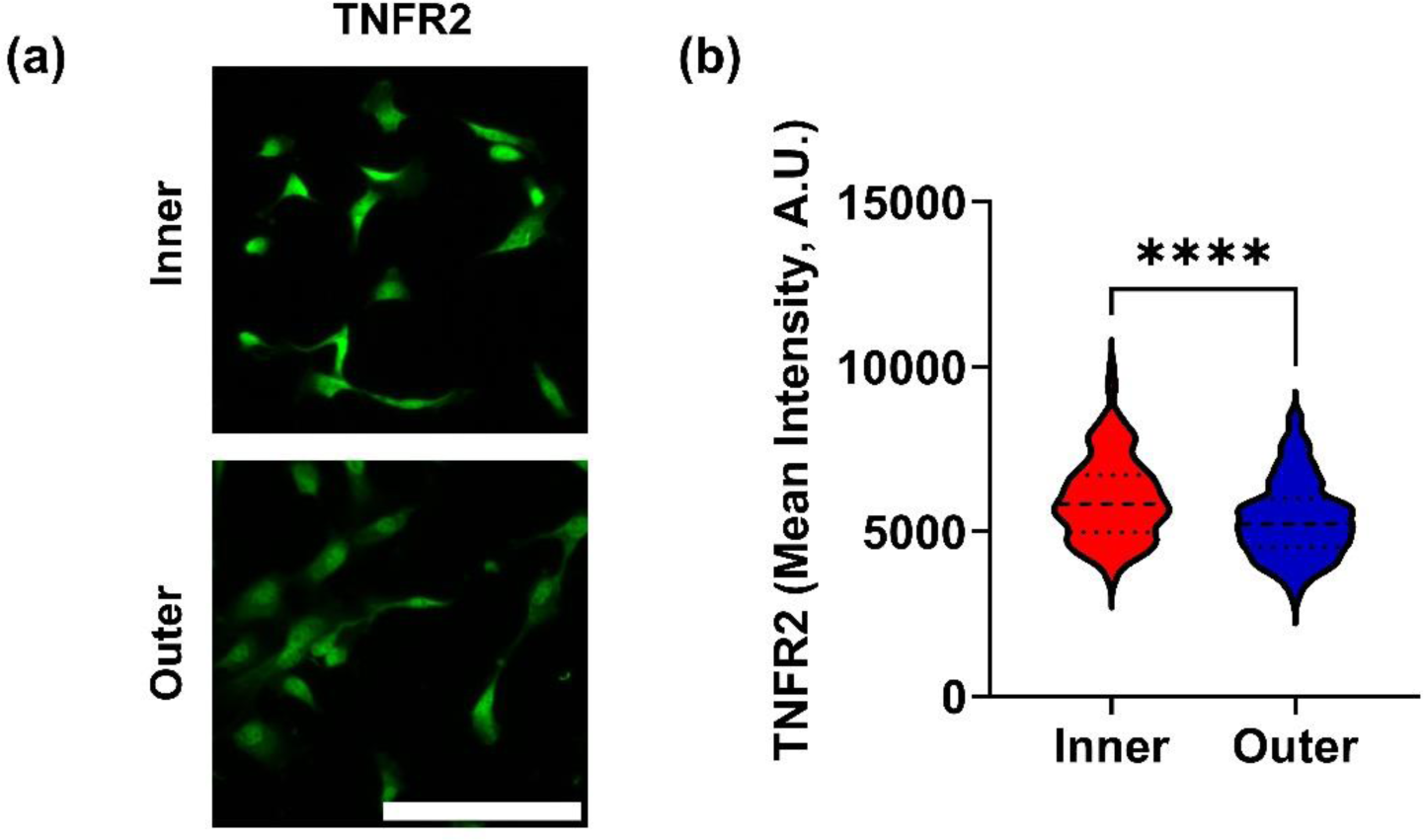
Baseline TNFR2 expressions in inner and outer meniscus cells. (a) Representative Immunofluorescence images of TNF-α Receptor 2 (TNFR2) in inner versus outer meniscus cells (scale bar = 100 μm). (b) Quantification of TNFR2 expression levels (n>100 cells/group, ****: p<0.0001, Mean +/-SD).

**Supplementary Table 1.**
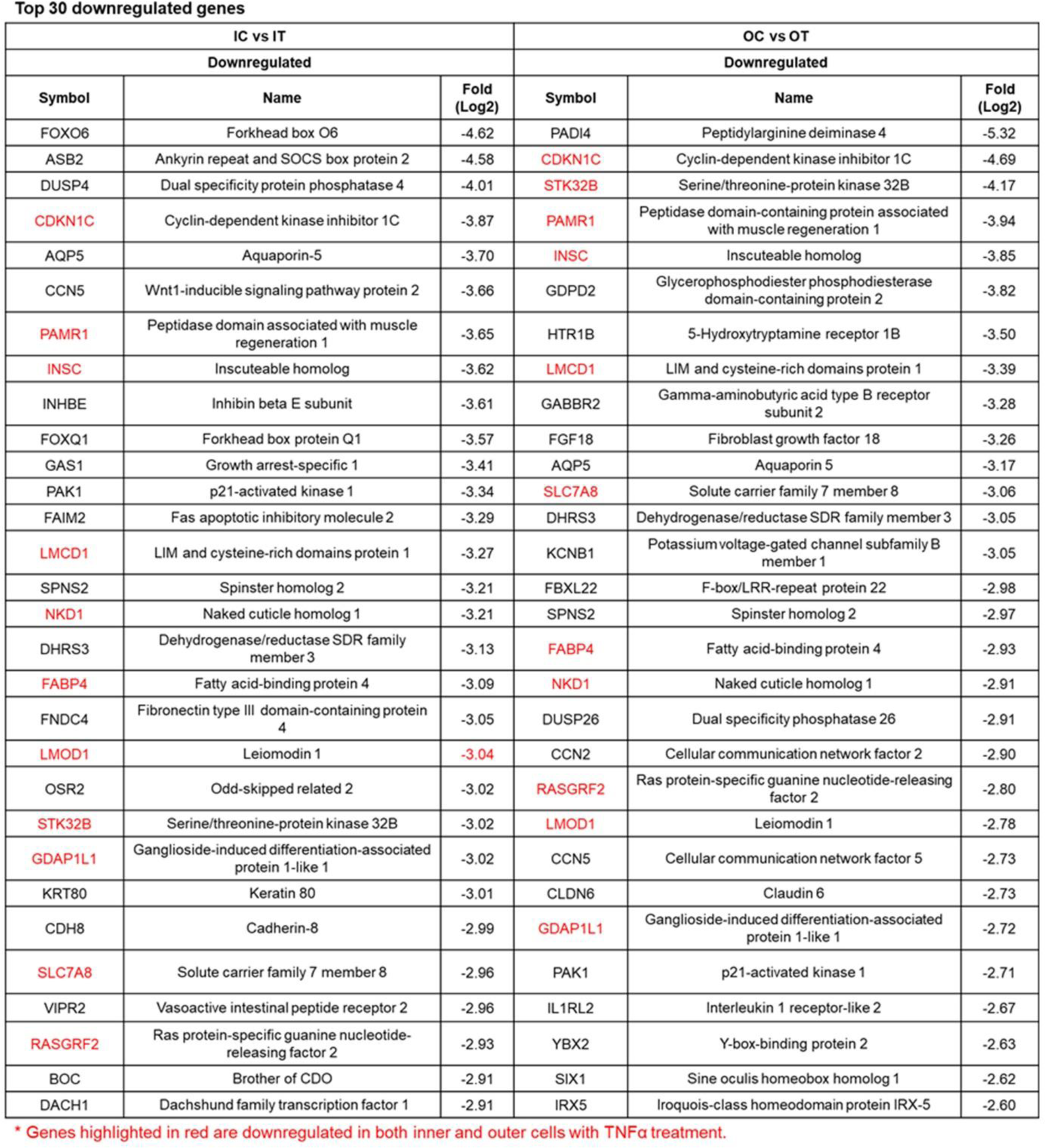
List of top 30 downregulated genes of inner and outer meniscus cells after TNF-α Treatment.

**Supplementary Table 2.**
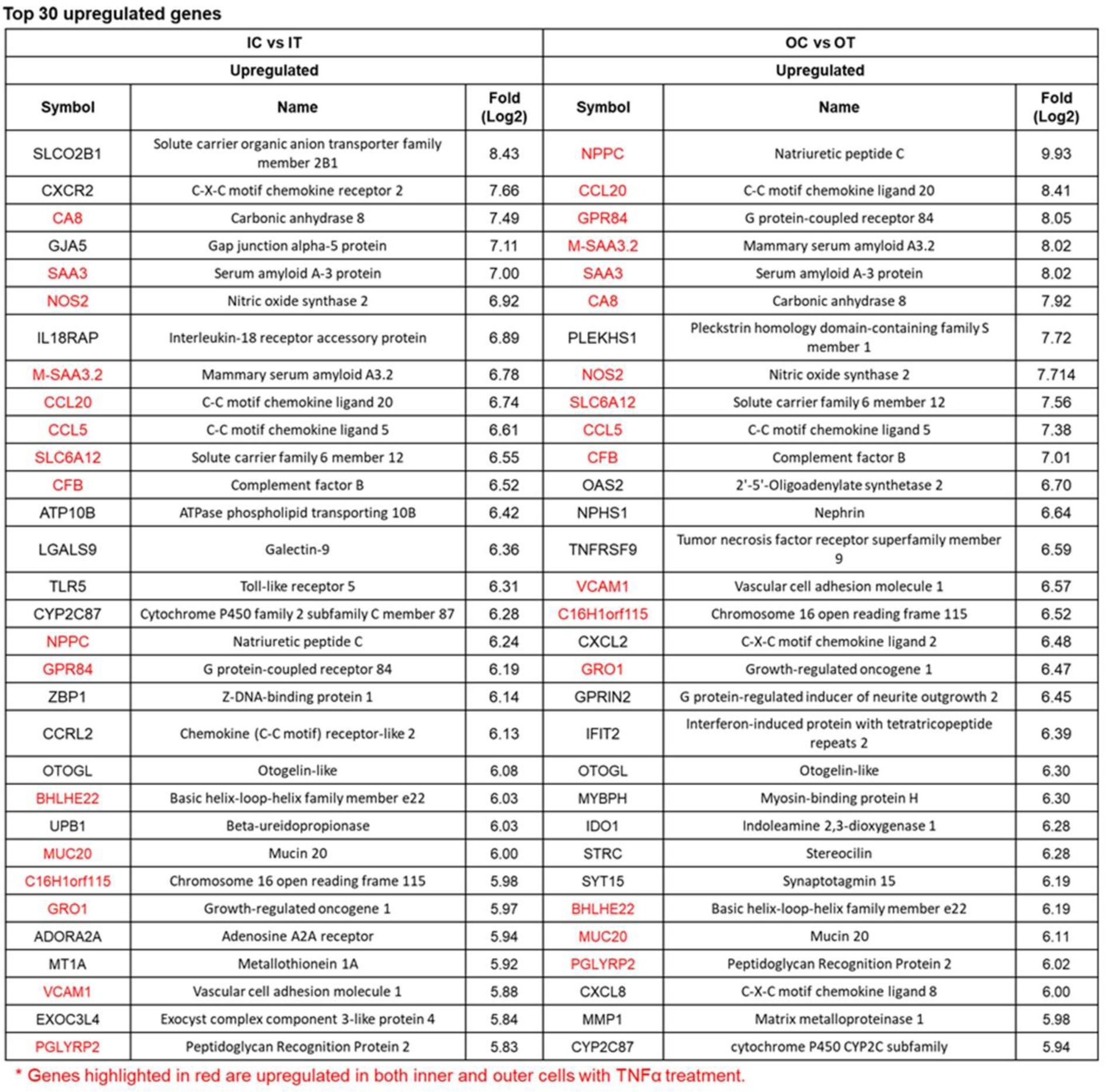
List of top 30 upregulated genes of inner and outer meniscus cells after TNF-α Treatment.

