## Supplementary information for "Epigenetic Dynamics in Meniscus Cell Migration and its Zonal Dependency in Response to Inflammatory Conditions: Implications for Regeneration Strategies"

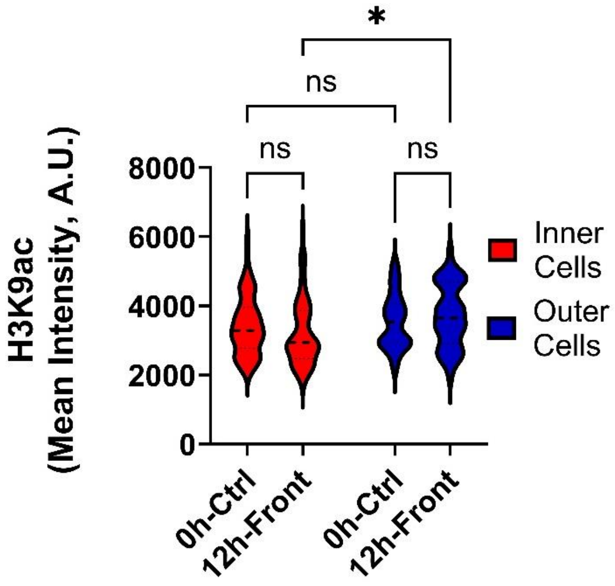


**Supplementary Figure 1: Changes in H3K9ac levels during inner and outer meniscus cell migration**. Quantification of H3K9ac Immunofluorescence results for Inner and Outer meniscus cells, measured before and 12 hours after scratch (n > 50 cells/group). *: p<0.05, Mean± SD).

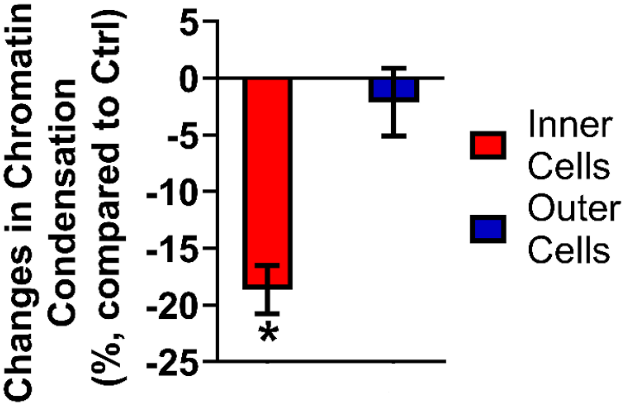


**Supplementary Figure 2: Global chromatin condensation changes for inner and outer meniscus cell migration under TNF-α treatment.** Percent change in chromatin condensation in nuclei at the migration "front" for inner and outer meniscus cells after 12-hour TNF-α (50 ug/ml, n = 7/group, *: p<0.0001. Mean± SD).

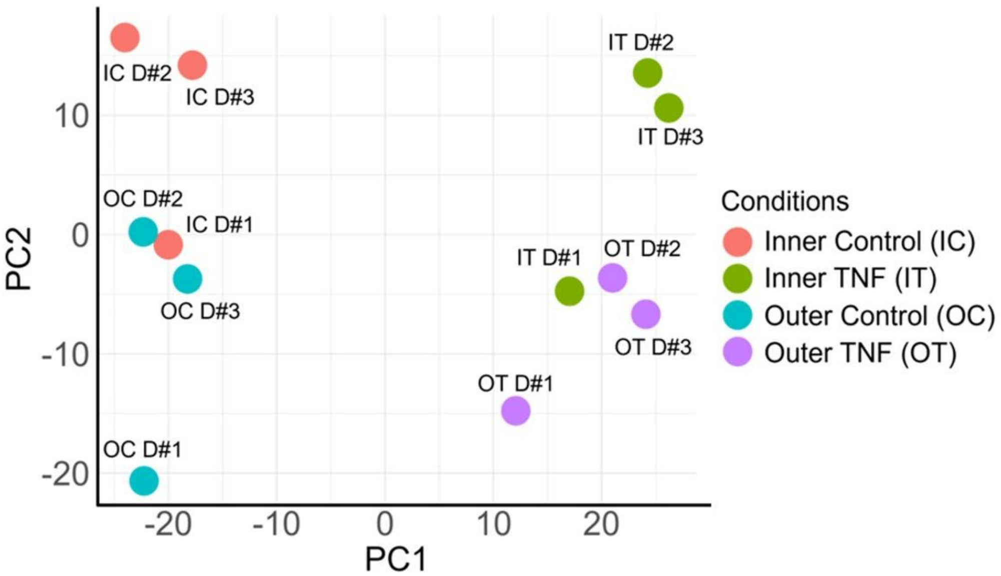


**Supplementary Figure 3**: **Global transcriptomic profiles of inner and outer meniscus cells.** Principal Component Analysis (PCA) plot illustrating the distinctions and similarities between the four experimental conditions based on their global gene expression profiles. Each point represents a sample, highlighting the clustering patterns that differentiate inner and outer meniscus cells under varying treatments.

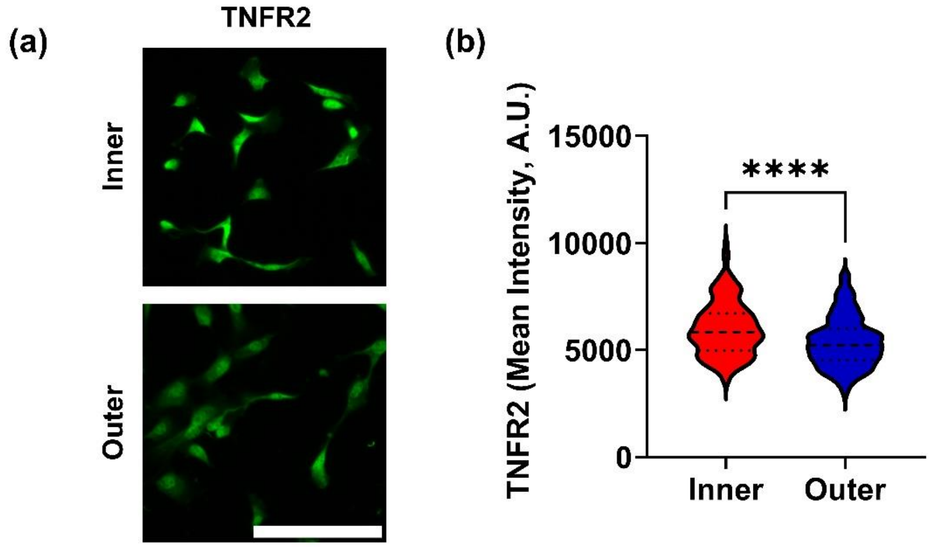


**Supplementary Figure 4: Baseline TNFR2 expressions in inner and outer meniscus cells.** (a) Representative Immunofluorescence images of TNF-α Receptor 2 (TNFR2) in inner versus outer meniscus cells (scale bar = 100 μm). (b) Quantification of TNFR2 expression levels (n>100 cells/group, ****: $p<0.0001$, Mean +/- SD).

**Supplementary Table 1**: **List of top 30 downregulated genes of inner and outer meniscus cells after TNF-α Treatment**

**Top 30 downregulated genes**

| IC vs IT | | | OC vs OT | | |
|---|---|---|---|---|---|
| Downregulated | | | Downregulated | | |
| Symbol | Name | Fold (Log2) | Symbol | Name | Fold (Log2) |
| FOXO6 | Forkhead box O6 | -4.62 | PADI4 | Peptidylarginine deiminase 4 | -5.32 |
| ASB2 | Ankyrin repeat and SOCS box protein 2 | -4.58 | CDKN1C | Cyclin-dependent kinase inhibitor 1C | -4.69 |
| DUSP4 | Dual specificity protein phosphatase 4 | -4.01 | STK32B | Serine/threonine-protein kinase 32B | -4.17 |
| CDKN1C | Cyclin-dependent kinase inhibitor 1C | -3.87 | PAMR1 | Peptidase domain-containing protein associated with muscle regeneration 1 | -3.94 |
| AQP5 | Aquaporin-5 | -3.70 | INSC | Inscuteable homolog | -3.85 |
| CCN5 | Wnt1-inducible signaling pathway protein 2 | -3.66 | GDPD2 | Glycerophosphodiester phosphodiesterase domain-containing protein 2 | -3.82 |
| PAMR1 | Peptidase domain associated with muscle regeneration 1 | -3.65 | HTR1B | 5-Hydroxytryptamine receptor 1B | -3.50 |
| INSC | Inscuteable homolog | -3.62 | LMCD1 | LIM and cysteine-rich domains protein 1 | -3.39 |
| INHBE | Inhibin beta E subunit | -3.61 | GABBR2 | Gamma-aminobutyric acid type B receptor subunit 2 | -3.28 |
| FOXQ1 | Forkhead box protein Q1 | -3.57 | FGF18 | Fibroblast growth factor 18 | -3.26 |
| GAS1 | Growth arrest-specific 1 | -3.41 | AQP5 | Aquaporin 5 | -3.17 |
| PAK1 | p21-activated kinase 1 | -3.34 | SLC7A8 | Solute carrier family 7 member 8 | -3.06 |
| FAIM2 | Fas apoptotic inhibitory molecule 2 | -3.29 | DHRS3 | Dehydrogenase/reductase SDR family member 3 | -3.05 |
| LMCD1 | LIM and cysteine-rich domains protein 1 | -3.27 | KCNB1 | Potassium voltage-gated channel subfamily B member 1 | -3.05 |
| SPNS2 | Spinster homolog 2 | -3.21 | FBXL22 | F-box/LRR-repeat protein 22 | -2.98 |
| NKD1 | Naked cuticle homolog 1 | -3.21 | SPNS2 | Spinster homolog 2 | -2.97 |
| DHRS3 | Dehydrogenase/reductase SDR family member 3 | -3.13 | FABP4 | Fatty acid-binding protein 4 | -2.93 |
| FABP4 | Fatty acid-binding protein 4 | -3.09 | NKD1 | Naked cuticle homolog 1 | -2.91 |
| FNDC4 | Fibronectin type III domain-containing protein 4 | -3.05 | DUSP26 | Dual specificity phosphatase 26 | -2.91 |
| LMOD1 | Leiomodin 1 | -3.04 | CCN2 | Cellular communication network factor 2 | -2.90 |
| OSR2 | Odd-skipped related 2 | -3.02 | RASGRF2 | Ras protein-specific guanine nucleotide-releasing factor 2 | -2.80 |
| STK32B | Serine/threonine-protein kinase 32B | -3.02 | LMOD1 | Leiomodin 1 | -2.78 |
| GDAP1L1 | Ganglioside-induced differentiation-associated protein 1-like 1 | -3.02 | CCN5 | Cellular communication network factor 5 | -2.73 |
| KRT80 | Keratin 80 | -3.01 | CLDN6 | Claudin 6 | -2.73 |
| CDH8 | Cadherin-8 | -2.99 | GDAP1L1 | Ganglioside-induced differentiation-associated protein 1-like 1 | -2.72 |
| SLC7A8 | Solute carrier family 7 member 8 | -2.96 | PAK1 | p21-activated kinase 1 | -2.71 |
| VIPR2 | Vasoactive intestinal peptide receptor 2 | -2.96 | IL1RL2 | Interleukin 1 receptor-like 2 | -2.67 |
| RASGRF2 | Ras protein-specific guanine nucleotide-releasing factor 2 | -2.93 | YBX2 | Y-box-binding protein 2 | -2.63 |
| BOC | Brother of CDO | -2.91 | SIX1 | Sine oculis homeobox homolog 1 | -2.62 |
| DACH1 | Dachshund family transcription factor 1 | -2.91 | IRX5 | Iroquois-class homeodomain protein IRX-5 | -2.60 |

* Genes highlighted in red are downregulated in both inner and outer cells with TNFα treatment.

**Supplementary Table 2**: **List of top 30 upregulated genes of inner and outer meniscus cells after TNF-α Treatment**

**Top 30 upregulated genes**

| IC vs IT | | | OC vs OT | | |
|---|---|---|---|---|---|
| Upregulated | | | Upregulated | | |
| Symbol | Name | Fold (Log2) | Symbol | Name | Fold (Log2) |
| SLCO2B1 | Solute carrier organic anion transporter family member 2B1 | 8.43 | NPPC | Natriuretic peptide C | 9.93 |
| CXCR2 | C-X-C motif chemokine receptor 2 | 7.66 | CCL20 | C-C motif chemokine ligand 20 | 8.41 |
| CA8 | Carbonic anhydrase 8 | 7.49 | GPR84 | G protein-coupled receptor 84 | 8.05 |
| GJA5 | Gap junction alpha-5 protein | 7.11 | M-SAA3.2 | Mammary serum amyloid A3.2 | 8.02 |
| SAA3 | Serum amyloid A-3 protein | 7.00 | SAA3 | Serum amyloid A-3 protein | 8.02 |
| NOS2 | Nitric oxide synthase 2 | 6.92 | CA8 | Carbonic anhydrase 8 | 7.92 |
| IL18RAP | Interleukin-18 receptor accessory protein | 6.89 | PLEKHS1 | Pleckstrin homology domain-containing family S member 1 | 7.72 |
| M-SAA3.2 | Mammary serum amyloid A3.2 | 6.78 | NOS2 | Nitric oxide synthase 2 | 7.714 |
| CCL20 | C-C motif chemokine ligand 20 | 6.74 | SLC6A12 | Solute carrier family 6 member 12 | 7.56 |
| CCL5 | C-C motif chemokine ligand 5 | 6.61 | CCL5 | C-C motif chemokine ligand 5 | 7.38 |
| SLC6A12 | Solute carrier family 6 member 12 | 6.55 | CFB | Complement factor B | 7.01 |
| CFB | Complement factor B | 6.52 | OAS2 | 2'-5'-Oligoadenylate synthetase 2 | 6.70 |
| ATP10B | ATPase phospholipid transporting 10B | 6.42 | NPHS1 | Nephrin | 6.64 |
| LGALS9 | Galectin-9 | 6.36 | TNFRSF9 | Tumor necrosis factor receptor superfamily member 9 | 6.59 |
| TLR5 | Toll-like receptor 5 | 6.31 | VCAM1 | Vascular cell adhesion molecule 1 | 6.57 |
| CYP2C87 | Cytochrome P450 family 2 subfamily C member 87 | 6.28 | C16H1orf115 | Chromosome 16 open reading frame 115 | 6.52 |
| NPPC | Natriuretic peptide C | 6.24 | CXCL2 | C-X-C motif chemokine ligand 2 | 6.48 |
| GPR84 | G protein-coupled receptor 84 | 6.19 | GRO1 | Growth-regulated oncogene 1 | 6.47 |
| ZBP1 | Z-DNA-binding protein 1 | 6.14 | GPRIN2 | G protein-regulated inducer of neurite outgrowth 2 | 6.45 |
| CCRL2 | Chemokine (C-C motif) receptor-like 2 | 6.13 | IFIT2 | Interferon-induced protein with tetratricopeptide repeats 2 | 6.39 |
| OTOGL | Otogelin-like | 6.08 | OTOGL | Otogelin-like | 6.30 |
| BHLHE22 | Basic helix-loop-helix family member e22 | 6.03 | MYBPH | Myosin-binding protein H | 6.30 |
| UPB1 | Beta-ureidopropionase | 6.03 | IDO1 | Indoleamine 2,3-dioxygenase 1 | 6.28 |
| MUC20 | Mucin 20 | 6.00 | STRC | Stereocilin | 6.28 |
| C16H1orf115 | Chromosome 16 open reading frame 115 | 5.98 | SYT15 | Synaptotagmin 15 | 6.19 |
| GRO1 | Growth-regulated oncogene 1 | 5.97 | BHLHE22 | Basic helix-loop-helix family member e22 | 6.19 |
| ADORA2A | Adenosine A2A receptor | 5.94 | MUC20 | Mucin 20 | 6.11 |
| MT1A | Metallothionein 1A | 5.92 | PGLYRP2 | Peptidoglycan Recognition Protein 2 | 6.02 |
| VCAM1 | Vascular cell adhesion molecule 1 | 5.88 | CXCL8 | C-X-C motif chemokine ligand 8 | 6.00 |
| EXOC3L4 | Exocyst complex component 3-like protein 4 | 5.84 | MMP1 | Matrix metalloproteinase 1 | 5.98 |
| PGLYRP2 | Peptidoglycan Recognition Protein 2 | 5.83 | CYP2C87 | cytochrome P450 CYP2C subfamily | 5.94 |

* Genes highlighted in red are upregulated in both inner and outer cells with TNFα treatment.
